# Cinnamon hydro alcoholic extract increased expression level of UCP3 gene in skeletal muscle of obese male Wistar Rats

**DOI:** 10.1101/105155

**Authors:** Elham Golmohamadi, Sanaz Mahmazi, Medi Rahnema

## Abstract

UCPs are the mitochondrial inner membrane proteins regulating basal metabolism. Ucp3 in muscle and adipose tissue helping metabolism and fat oxidation affecting thermogenesis, involve in fatty acid metabolism and energy homeostasis.

42 adult rats divided into 6 groups including control group, the high-fat diet Obese group, experimental groups that in addition to the high-fat diet received 50,100,200 mgKg^-1^ doses of cinnamon extract and the sham group with 200 mgKg^-1^ extract treatment. RNA extracted from muscles, cDNA synthesized and Ucp3 gene expression level was examined by real-time PCR.

In Obese rats’ muscle, significant decrease in UCP3 gene expression was observed but in exprimental groups there was significant increased level of UCP3 gene expression in 100, 50 mgKg^-1^ dose of treatment.

UCP3 might be a target for pharmacological up regulation in treatment of obesity. Cinnamon might influence UCP3 expression, in a dose dependent manner and low dose was more effective.

## Introduction

Obesity has become a significant clinical problem in recent decades since it could contribute to life-threatening diseases. It is characterized by increased fat mass, which is associated with increased cell number and sizes. Adipogenesis and fat accumulation play an important role in fat mass increase (Lee et al., 2015). Childhood Obesity could significantly increases the risk of hypertension, dyslipidemia and type 2 diabetes (Ghibaudi et al., 2002). The model of diet-induced obesity has many features like high fat diet (Levin and Dunn-Meynell, 2006).

Any factors that could decreased Fatty Acid level and enhance their metabolism or increase lipolysis and prevent lipogenesis could be a determinative factor for obesity prevention and control. Some foods and herbs by interference metabolism could facilitate fatty acid oxidation and prevent obesity (Chu et al., 2015, Ali et al., 2015).

In traditional medicine cinnamon (***Cinnamomum zeylanicum***) or Dalchini is a herb used for gastrointestinal problems, urinary infections, relieving symptoms of colds and flu and has remarkable anti-bacterial properties. Some studies have shown that Cinnamon reducing fasting blood glucose and systolic blood pressure, and improving body composition in men and women with the metabolic syndrome and can reduce risk factors associated with diabetes, Obesity and cardiovascular diseases (Askari et al., 2014).

Researches shown that cinnamon bark powder at different doses 1, 3 and 6 g/day could prevents hypercholesterolemia and hypertriglyceridemia and lowers the levels of free fatty acids and triglycerides in plasma of type 2 diabetic subjects by its strong lipolytic activity (Askari et al., 2014, Khan et al., 2003). Cinnamate, a phenolic compound found in cinnamon bark and other plant materials, lowers cholesterol levels in high fat-fed rats by inhibiting hepatic 5-hydroxy-3- methylglutaryl-coenzyme A (HMG-CoA) reductase activity (Kannappan et al., 2006).

Thermogenesis has been implicated in the regulation of body temperature, body weight and metabolism. Mitochondrial proteins called uncoupling proteins (UCPs) induce non shivering thermogenesis by creating a pathway that allows dissipation of the proton electrochemical gradient across the inner mitochondrial membrane (Matsuda et al., 1997). UCP3 plays critical role in fatty acid metabolism and anions transport. It was shown that UCP1 and UCP3 expression contributed to lipid and carbohydrate oxidation, UCP3 expression was associated with BMI and percentage of lean body mass in human (Oliveira et al., 2016). UCP3 expression in skeletal muscle and brown adipose tissue regulate energy consumption by fatty acid oxidation and has been associated with whole-body energy metabolism. The other function of UCP3 was involvement in the regulation of reactive oxygen species production, mitochondrial fatty acid transport and glucose metabolism regulation in skeletal muscle (Schrauwen, 2002).

UCP3 has an important role in fatty acid anions transportation from mitochondrial matrix before oxidation, thereby protecting mitochondria against lipid-induced damage (Schrauwen, 2004). UCP2 and UCP3 as mitochondrial electron transporters are involved in ATP production and energy wastage regulation as heat. Energy output has an important role in physical performance, especially in aerobic fitness (Holdys et al., 2013). UCP3 could be an interesting target for pharmacological up regulation in the treatment of obesity and diabetes. Then in this study we aimed to investigate how cinnamon extract influences UCP3 gene expression in hand Epitroch Brachii and foot Gracilis muscles of high fat dietary obese rats.

## Materials and methods

## Rats and Diets

Male Wistar rats (n = 42) were obtained from The Zanjan University of Medical Sciences and individually housed at the Islamic Azad University of Zanjan’s Animal House. All experimental protocols were approved by the Animal Experimentation Ethics Committee of the Zanjan University of Medical Sciences, under the guidelines of the National Health and Medical Research Council of Australia. 42 male Wistar rats were randomly divided into 6 groups: The control group with a normal diet, three experimental groups received high-fat diet and Cinnamon 70% hydro alcoholic extract in 50 mgKg^-1^, 100 mgKg^-1^ and 200 mgKg^-1^ doses. Obese group received high-fat diet and Sham group received normal diet and 200 mgKg^-1^ cinnamon extract. All groups treated for 6 weeks with oral gavage of high-fat emulsion as 1.5 ml (table A1) once per day and Cinnamon hydro alcoholic extract. The rats were individually housed in a temperature-controlled room (22°C) under a 12 h light– 12 h dark cycle. At the end of the experimental period, the rats were fasted for 12 h, anesthetized with Somnopentyl (pentobarbital sodium 64.8 mg/mL solution). The muscle tissues were collected, weighed and immediately used for RNA extraction.

Cinnamon 70% Hydroalcoholic extract was ordered from Iranian biological resource center, Karaj, Iran.

**Table A1:** The composition of high-fat emulation

| components | Weight in emulsion |
| --- | --- |
| corn oil | 400 g |
| Sucrose | 150 g |
| milk powder | 80 g |
| Cholesterol | 100 g |
| Tween 80 | 36.4 g |
| Propylene glycol | 31.1 g |
| multi vitamine | 2.5 g |
| Salt | 10 g |
| Distilled water | 300 g |

## Gene expression

Total RNA was isolated from muscle tissue using RNX kit (Cinnagene). Messenger RNA was reverse transcribed into cDNA using a Takara cDNA Synthesis kit (Takara). Q real-time PCR was used to validate a subset of differentially expressed genes. Gene-specific primers were designed using the Primer3 program (http://frodo.wi.mit.edu/). Primer design criteria included a base-pair length of 100 to 200 and a guanine-cytosine (GC) content of 40% to 60%. Primers were designed to span exon/intron boundaries where possible and were tested using the same cDNA sample pool, to ensure that there was no genomic contamination. We used UCP3 f 5`- GACTCACAGGCAGCAAAGGAA- 3` and UCP3 r 5`-GAGGAGATCAGCAAAACAGGC-3` specific primers that produced 131bp products and B2M primers F 5`- CGAGACCGATGTATATGCTTGC-3` and R 5`-GTCCAGATGATTCAGAGCTCCA-3` as refrence gene specific primers. Real-time PCR analysis was performed using SYBR Green (Ampliqon- Danmark). Each 10 μL reaction contained 2X SYBR Green Master Mix I, 0.5 μmol/L gene specific forward and reverse primers, and 100 ng cDNA. Polymerase chain reaction analysis was accomplished using the Rotor-Gene Q PCR System with the following cycle conditions: 95°C for 15 minutes followed by 40cycles of 95°C for 10 seconds, 60°C for 30 seconds, and 72°C for 30 seconds. A melt curve analysis confirmed the amplification of a single cDNA product. Relative quantification of UCP3, toward the samples, were calculated by using the comparative2^-△△Ct^ method. B2M is the most stable reference genes for muscle tissue.

## Statistical analysis

Descriptive statistics consisted of mean values and standard deviation. Data normality was verified by the T-test. Statistical significance was set at (p < 0.05).

## Result and Discussion

In this study high fat diet fed obese group rats had significant weight gain. We demonstrated that cinnamon extract significantly decreased body mass index BMI in sham group but in experimental groups compared to obese group only 100(3.827±0.256) and 50(3.791±0.351) mgKg^-1^ dose of cinnamon extract decreased BMI significantly (P<0.05). Cinnamon extract in high dose 200 mgKg^-1^ (4.895±0.297) with high fat diet could not decrease body weight and BMI (fig-A1).

**Fig A1:**
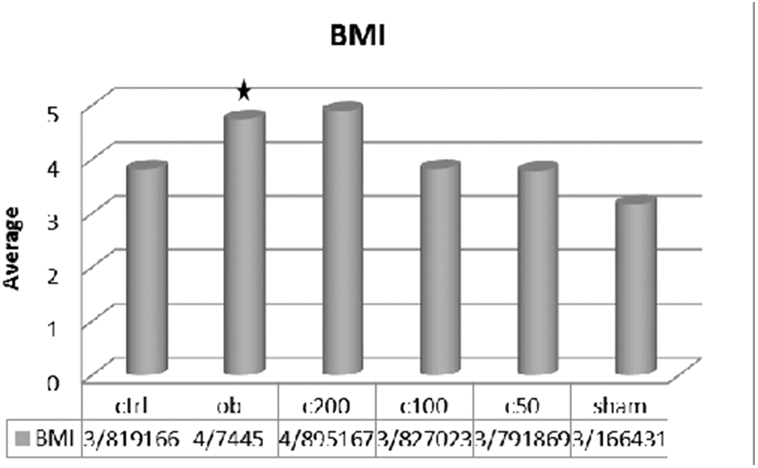
Body mass index in studied groupes

UCP3 gene expression was evaluated in leg (Gracilis muscle) and hand (epitroch brachii muscle) muscles of all studied groups. In leg muscle of obese group there was significant decreased in UCP3 gene expression (0.1340±0.189) (P<0.05) but in all experimental groups with high fat diet and 50(23.479± 10.965), 100(11.1062±6.225) and 200(2.2641±0.661) mgKg^-1^ cinnamon extract treatment UCP3 mRNA level significantly increased (P<0.05). In sham group with 200 mgKg^-1^ cinnamon extract treatment there was not significant increase (P>0.05). We didn’t investigate significant differences in UCP3 gene expression level in hand muscle of all studied groups (fig-A2). As a results cinnamon extract in low dose and in leg skeletal muscle could increase UCP3 gene expression.

**Fig A2:**
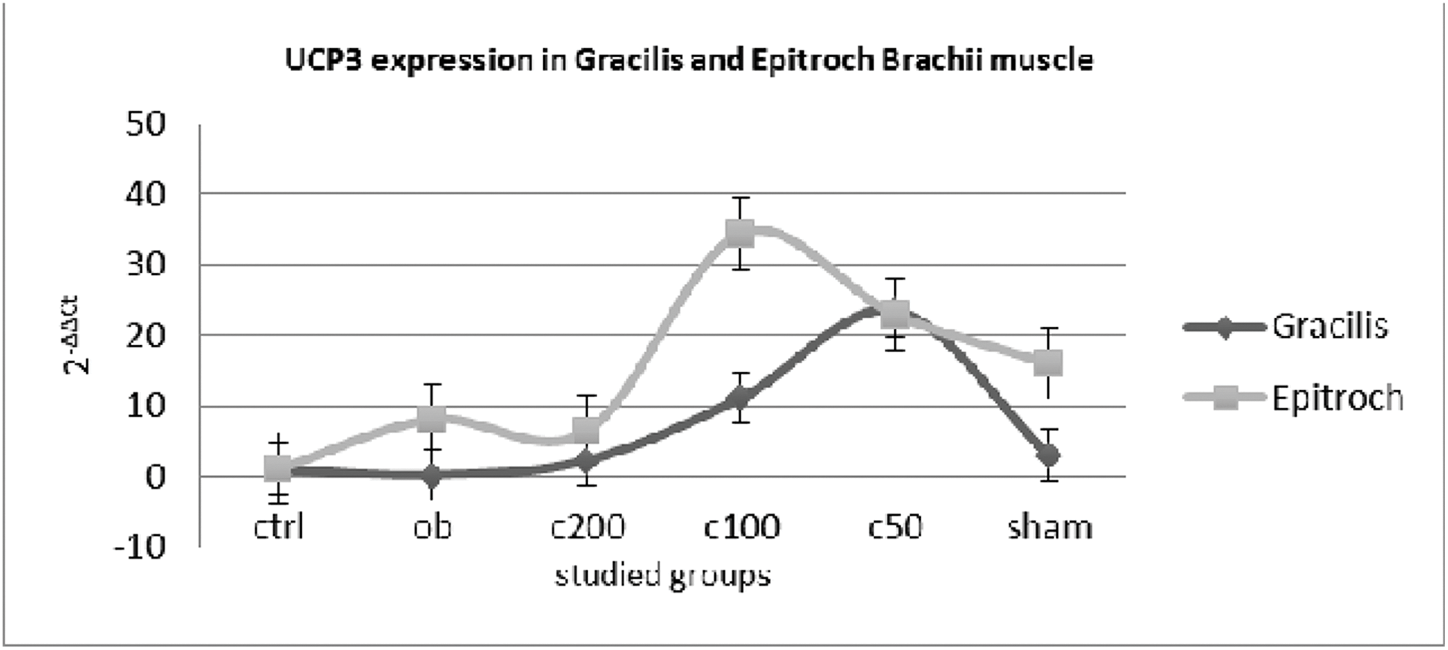
UCP3 gene expression fold changes in Gracilis and Epitroch muscles of studied groups.

UCP3 is expressed predominantly in skeletal muscle and has been associated with whole-body energy metabolism. In mitochondrial regulates fatty acid transport and in skeletal muscle, regulates glucose metabolism (Oliveira et al., 2016, Schrauwen, 2004). Study of diet-resistant obese women shown that expression of UCP3 mRNA abundance in skeletal muscle was 25% greater in diet-responsive than in diet-resistant subjects. Thus, proton leak and the expression of UCP3 correlate with weight loss success and may be candidates for pharmacological regulation of fat oxidation in obese diet-resistant subjects (Harper et al., 2002). As our found the *Cinnamon zeylanicum* hydro alcoholic extract could increase UCP3 gene expression in high fat dietary obese Rats and reduced their weight and BMI then cinnamon extract could be a good supplement for obesity prevention and in diet resistant condition it might be helpful.

For analysis of UCP3 gene expression level we used hand Epitroch Brachii muscle and foot Gracilis muscle because of physiological and metabolic differences in skeletal muscles. The metabolism of mitochondria isolated from functionally distinct skeletal muscles is different (Rasmussen et al., 2004). In mammalian and avian Skeletal muscle fuel selection occurs at the mitochondrial level. different muscles according their structure and function might be have different number of mitochondria and use different sources of fuels like lipids and glucose (Kuzmiak-Glancy and Willis, 2014). As a different structure and function of Epitroch and Gracilis skeletal muscles and similarity of these muscles in human and rat we examine the effect of cinnamon extract on UCP3 expression level in those different muscles. In Epitroch muscle there wasn’t significant differences in all studied groups but in Gracilis muscle high fat diet and Obesity lead to decreased UCP3 expression. In experimental groups that received 50 and 100 mgKg^-1^ dose of cinnamon extract significant increased were seen. Cinnamon extract as an economic readily available food supplement could be used as an affective anti-obesity herb (Anderson et al., 2015).

Cinnamon extract might be decreased high fat diet effects and prevented obesity in high fat diet fed rats. High dose of Sinnamon was not effective and could not decreased body weight after high fat dietary treatment it shown that Cinnamon act dose dependent.

## Acknowledgements

This study was conducted in molecular genetic laboratory of Azad University, Zanjan Branch, Zanjan, Iran. We thanks A. Nazari for experimental assistance.

## Author contributions

S.M. and M.R. designed the study, M.R. and E.G. performed animal model experiments.

E.G. and S.M. performed molecular genetic experiments and analyzed data, S.M., and E.G. wrote the paper.

